## Supplementary Information for "Precision dynamical mapping using topological data analysis reveals a unique hub-like *transition state* at rest"

### Supplementary Figures

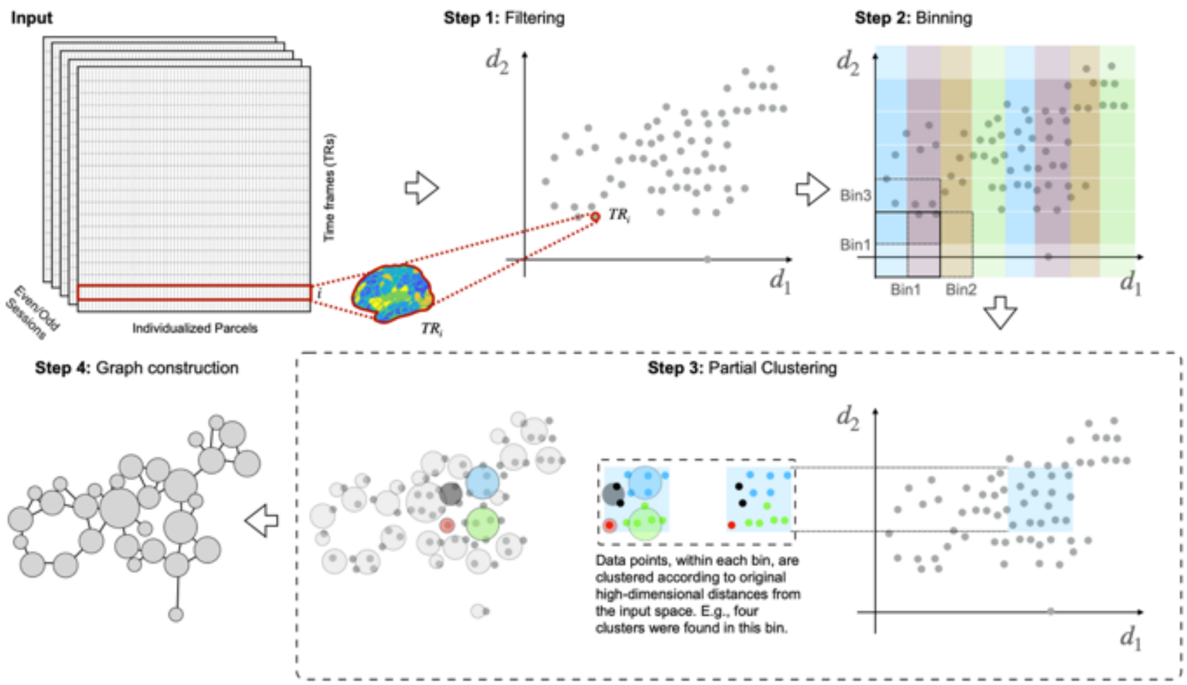


**Fig. S1: Step-by-step representation of the Mapper pipeline.** First, the high-dimensional neuroimaging data are embedded into a lower dimension set $d_i$, using a non-linear filter function $f$. Here, a nonlinear filter function $f$ based on neighborhood embedding was used (see Methods for benefits of this non-linear approach). Second, overlapping $d$-dimensional binning is performed to allow for compression and to reduce the destructive effects of noise. Third, partial clustering within each bin is performed, where the original high dimensional information is used for coalescing (or separating) data points into nodes in the low-dimensional space and hence allows for recovering information loss incurred due to dimensional reduction. As a fourth step, to generate a graphical representation of the data landscape, nodes from different bins are connected if any data points are shared between them.

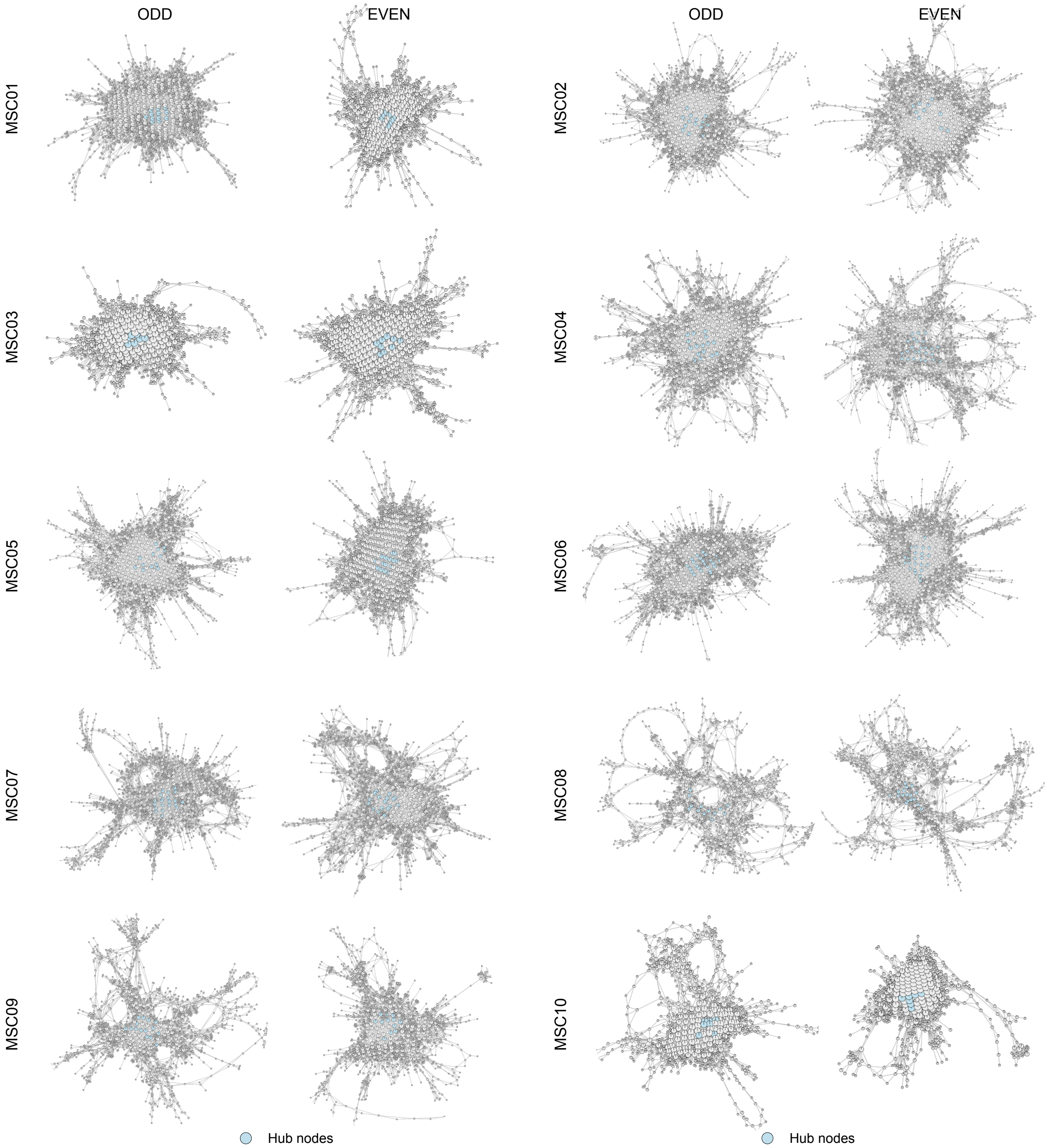


**Fig. S2: Shows hub nodes across all ten MSC participants.** Show Mapper-generated graphs for all 10 MSC participants and their respective data splits. Hub nodes (i.e., nodes with high degree (>20) and high centrality (top 1%)) are highlighted in blue color. As evident, these hub nodes were found across all participants and sessions.

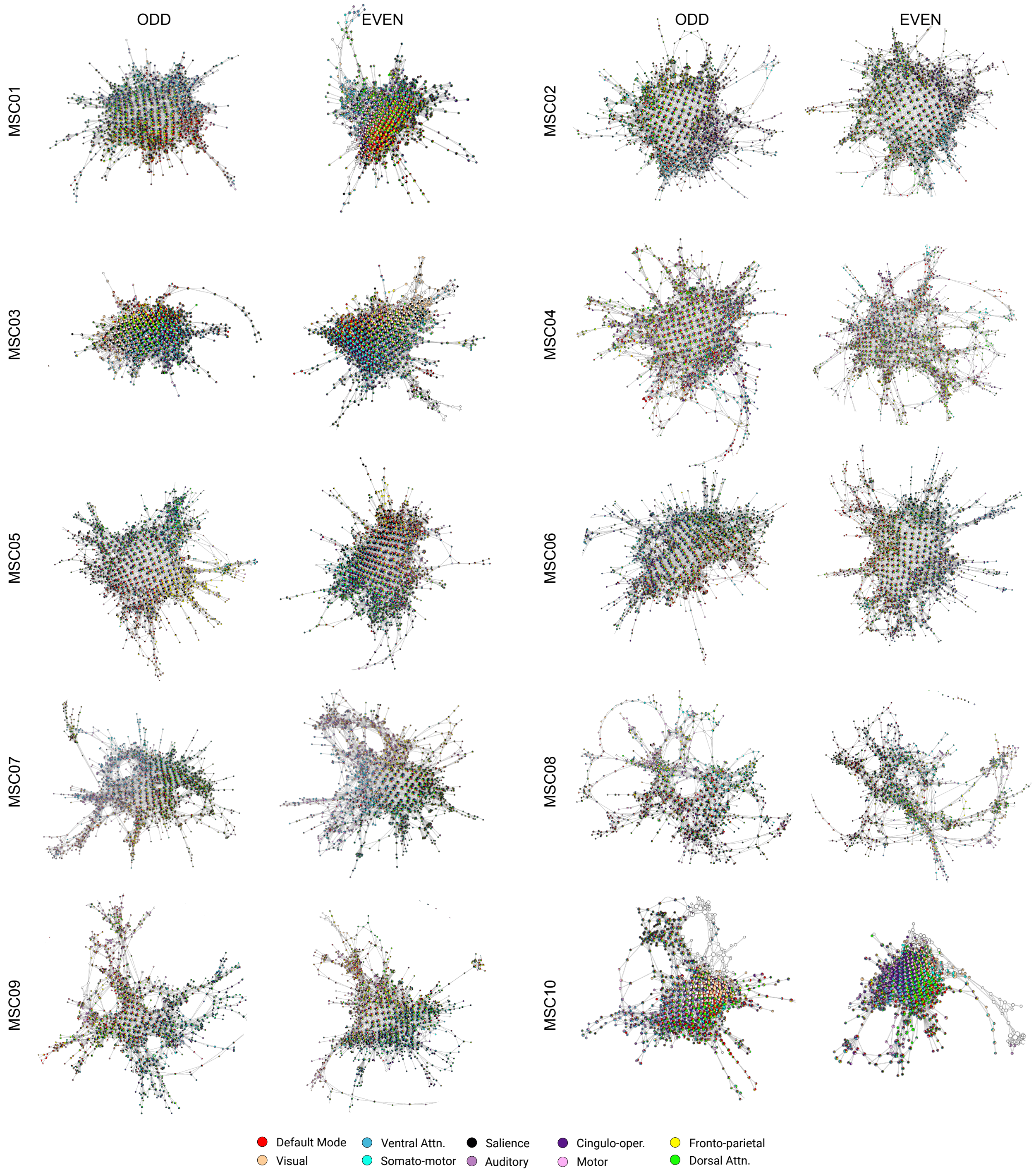


**Fig. S3: RSN-based annotation across all ten MSC participants.** Shows Mapper-generated graphs for all 10 MSC participants. Here, each node is annotated by activation in the known large-scale resting state networks. Each node is annotated using a pie-chart to show the proportion of RSNs activated within each node. As evident, topologically highly connected, and central hub nodes contained brain volumes where no characteristic RSN was activated above the mean, whereas nodes with brain volumes dominating from one (or more) RSN(s) tend to occupy the peripheral corners of the landscape. The maps for all individual subjects demonstrated this same basic pattern, although there was evidence to suggest that different combinations of RSNs were dominant in different individuals.

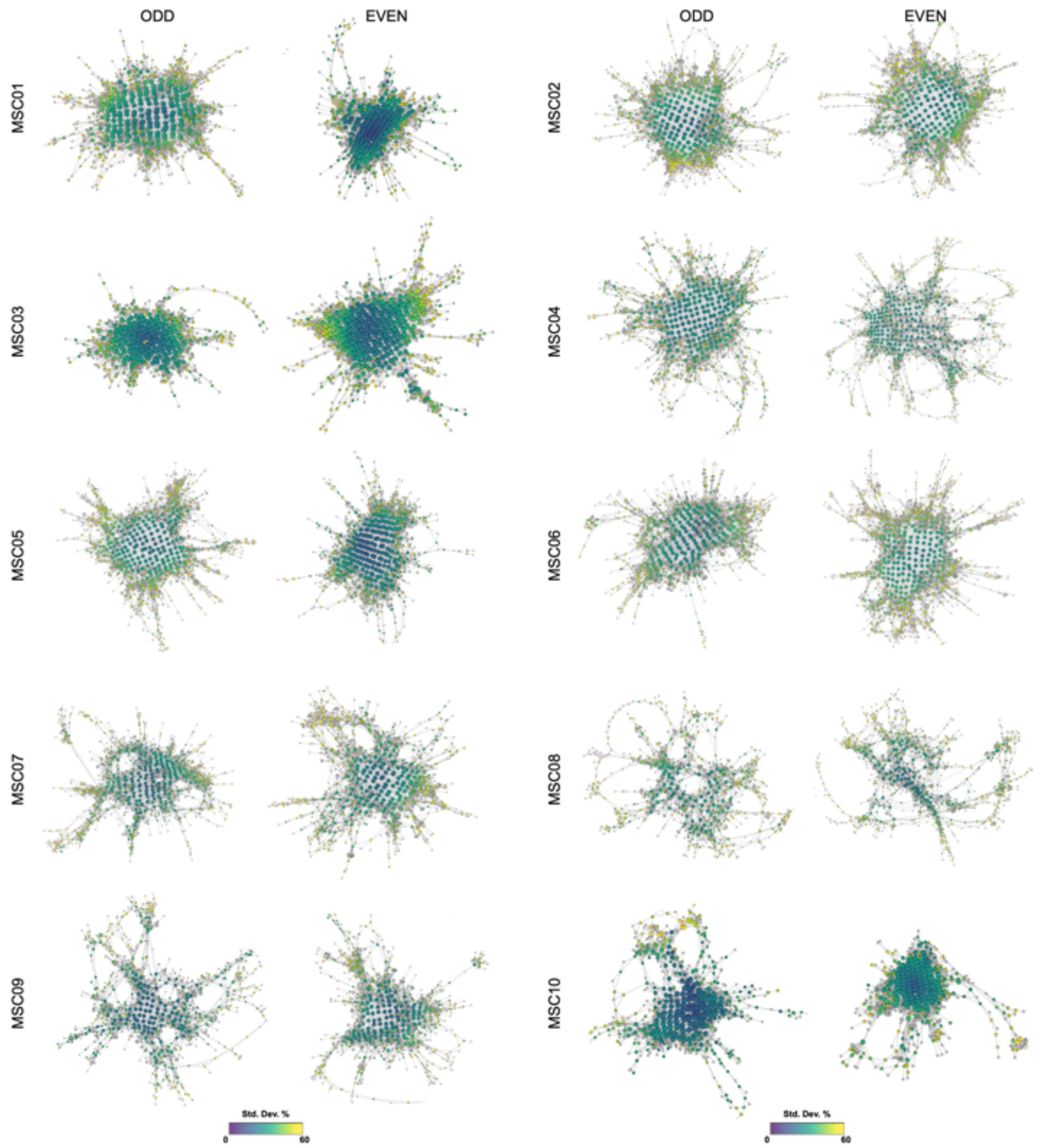


**Fig. S4: Gradient was observed across all ten MSC participants**. To quantify the variation in RSN-based dominance, we first estimated mean activation for each RSN across the time frames within each node, followed by estimating variation in mean activation across RSNs. High variance (or S.D.) indicated dominance of one or more RSN while low variance (or S.D.) indicated uniformity across RSN activation. Annotating Mapper-generated graphs using variance-based approach revealed a dynamical topographic gradient, where the peripheral nodes had higher variance with a continual decrease in variance when going towards the center of the graph. This topographic gradient was observed across all participants and sessions.

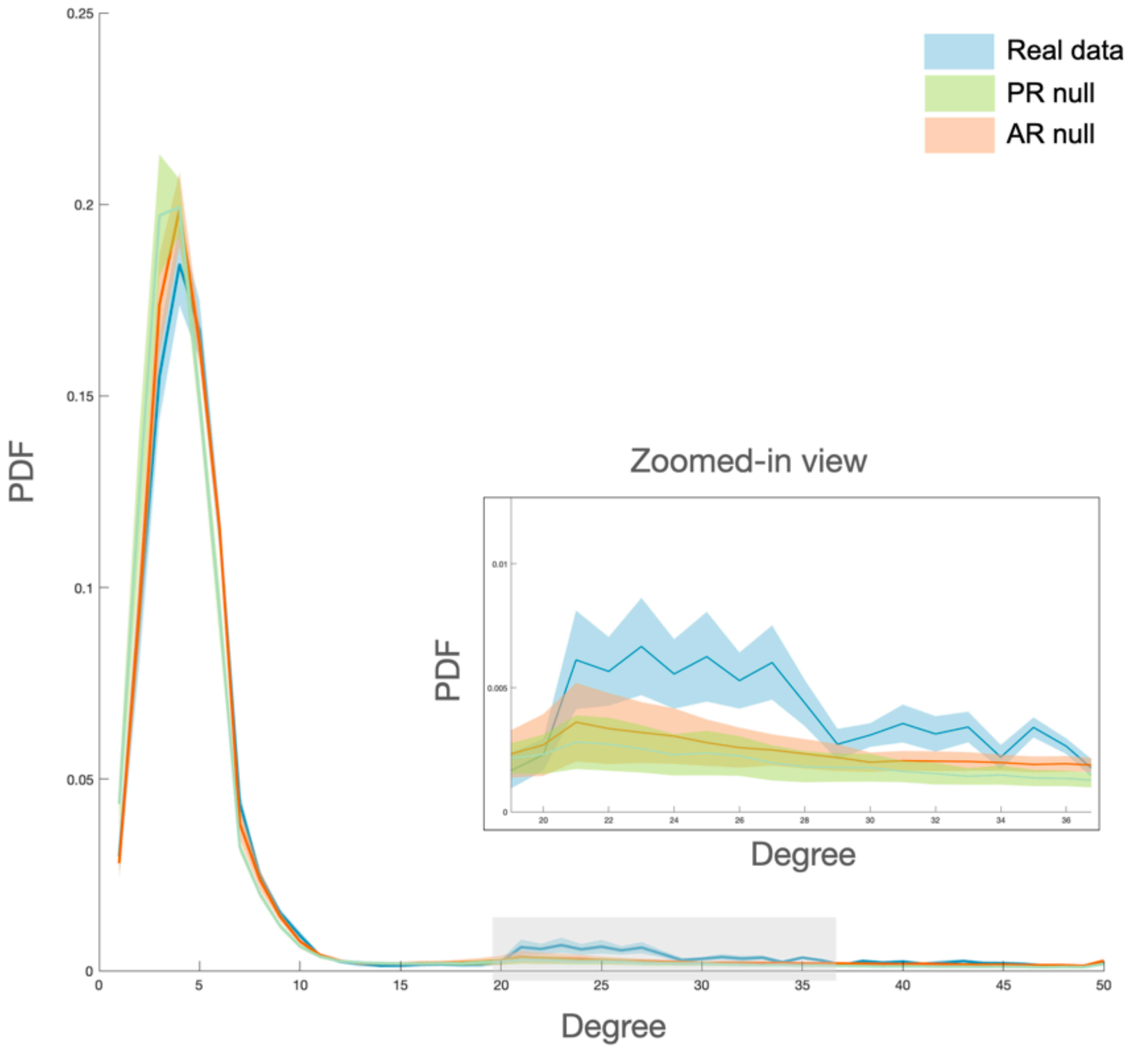


**Fig. S5: Applying frame censoring to the data generated from null models to evaluate whether high degree nodes observed in the real data were resulted merely due to temporal masking**. Please note that the frame censoring could only be applied to the data generated from the AR null, as it is a generative model, and we could create same number of TRs as the original (non-censored) data and then drop frames from it to match the number of frames in the real data. Here, we kept the PR null (without frame censoring) for comparison. As evident, adding frame censoring to the data generated using AR null did not result in enhancing the number of high degree nodes. Statistically, the proportion of high degree nodes in real data were still significantly higher than both nulls ($F=6.15$, $p=0.0063$). Only showing data from the odd split, similar results were observed for the even split of the data.

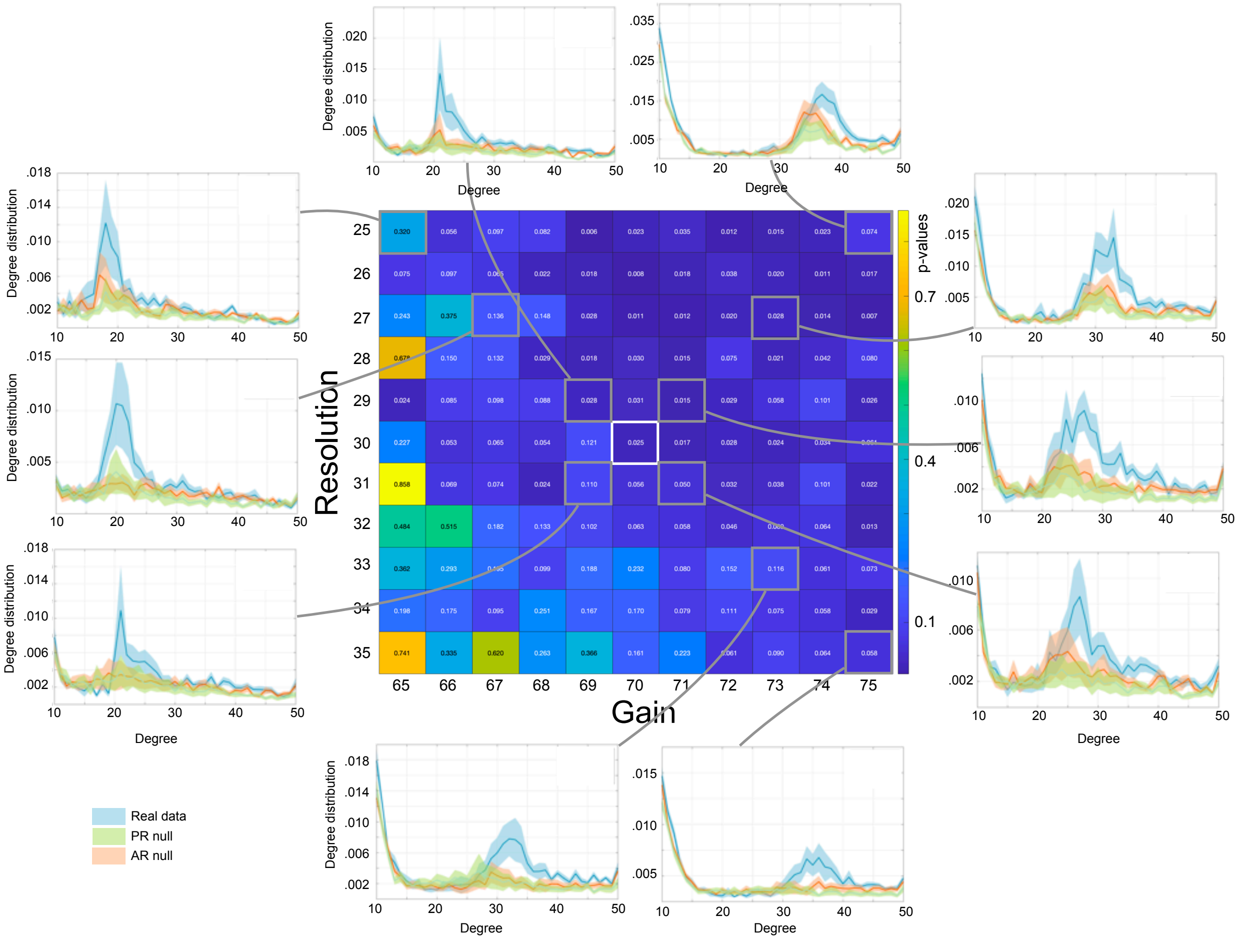


**Fig. S6: Parameter perturbation analysis revealed stable results across a wide range of Mapper parameters.** Parameter perturbation analysis was performed to make sure topological properties of the graph (e.g., existence of fat tail in degree distribution) were stable across a moderate range of Mapper parameters. Two main Mapper parameters, i.e., number of bins (a.k.a. Resolution (R)) and percentage overlap between bins (a.k.a. Gain (g)) were varied across the chosen value in the main text (R=30, G=70). The values of R and G were chosen based on our previous work with task fMRI data (Saggar et al. 2018; Nat. Comm.). The heatmap above shows p-values from one-way ANOVAs that examined the proportion of high-degree nodes (>20) in the real versus null data for the odd sessions. As evident, for a large portion of Mapper parameter values, the proportion of high-degree nodes in the real data were significantly higher than null data. We also depict zoomed-in view of degree distribution plots for several parameter combinations (highlighted in gray box on the heatmap) to show the excessive proportion of high-degree nodes in real data across different combinations of parameters.
